# Gating the channel pore of ionotropic glutamate receptors with bacterial substrate binding proteins

**DOI:** 10.1101/2021.01.27.428399

**Authors:** Max Bernhard, Bodo Laube

## Abstract

Tetrameric ionotropic glutamate receptors (iGluRs) mediate excitatory neurotransmission in the mammalian central nervous system and are involved in learning, memory formation, and pathological processes. Based on structural and sequence similarities of the ligand-binding and channel domains of iGluR subunits to bacterial binding proteins and potassium channels, iGluRs are thought to have originally arisen from their fusion. Here we report the functional coupling of the bacterial ectoine binding protein EhuB to the channel pore-forming transmembrane domains of the bacterial GluR0 receptor by stabilization of dimeric binding domains. Insertion of a disulfide bridge in the dimer interface abolished desensitization of the channel current analogous to mammalian iGluRs. These results demonstrate the functional compatibility of bacterial binding proteins to the gate of the channel pore of an iGluR. Moreover, our results highlight the modular structure and crucial role of binding domain dimerization in the functional evolution of iGluRs.

## Introduction

Tetrameric ionotropic glutamate receptors (iGluR) mediate the majority of excitatory neurotransmission in the central nervous system by converting a chemical signal (neurotransmitter) into an electrical signal (Traynelis et al., 2010). Based on their pharmacological properties, vertebrate iGluRs can be subdivided into four subfamilies: α-amino-3-hydroxy-5-methyl-4-isoaxazolepropionic acid (AMPA), kainate (KA), N-methyl-D-aspartate (NMDA), and δ-receptors. Additional members of the iGluR family can be found among the entire animal kingdom (Ramos-Vicente et al., 2018), in plants (Lam et al., 1998) and bacteria (Chen et al., 1999). Despite their diverse physiological functions, all eukaryotic iGluRs share the same modular architecture (Fig 1a). They consist of an extracellular N-terminal domain (NTD), an extracellular ligand-binding domain (LBD), a transmembrane domain (TMD; M1, M3, M4) including the pore loop (M2) and an intracellular C-terminal domain (CTD). The LBD is a two-lobed domain composed of an extracellular region (called D1 or S1) preceding the first transmembrane domain M1 and a second extracellular region (called D2 or S2) connecting the transmembrane segments M3 and M4. Glutamate occupation of the LBDs, which are arranged as two dimers in a “back-to-back” fashion in the tetrameric receptor, induces a closure of both clamshell-like domains that is transduced towards the TMD and results in permeation of Na^+^, K^+^ and Ca^2+^ - ions across the cell membrane (Armstrong et al., 2006; Dürr et al., 2014; Sobolevsky et al., 2009a). Nevertheless, the emergence of their modular architecture remains most widely unknown. The shared complex architecture of eukaryotic iGluRs suggests an ancient separation of the protein family (Lam et al., 1998; Price et al., 2012) with a common ancestor that can be traced back as far as bacteria (Chen et al., 1999; Janovjak et al., 2011). Prokaryotic iGluR subunits, such as GluR0 (Chen et al., 1999), display a simplified architecture, lacking the third transmembrane segment M4 and an NTD, while also exhibiting unique functional and pharmacological features, including a potassium selectivity filter and the ability to be activated by a broad range of amino acids. Interestingly, individual domain segments of iGluRs share structural similarities with other prokaryotic protein families. Thus, the LBD of iGluRs are structural homologue to class II bacterial solute binding proteins (SBP) (Felder et al., 1999) and the TMD displays an inverted architecture of tetrameric K^+^-channels (Doyle et al., 1998; Sobolevsky et al., 2009b), leading to the proposal that iGluRs possibly arose by the fusion of SBP and potassium channels. Recent findings confirm the general kinship between K^+^-channels and iGluRs by coupling an iGluR LBD to a small viral K^+^-channel (Schönrock et al., 2019), highlighting a conserved activation mechanism of the channel pore in both protein families. However, the overall sequence identities between iGluRs and SBPs are low (Tikhonov and Magazanik, 2009) and the evolutionary link between SBP and iGluR remains unclear. SBP mediate the uptake of a large variety of substances across the cell membrane (Tam and Saier, 1993) and is further involved in chemotaxis and DNA regulation (Lewis et al., 1996). Despite a low sequence similarity, SBP display, like iGluR LBDs, a highly conserved three-dimensional clamshell-like structure composed of two α/β domains (D1 and D2), connected by a small hinge with one to three interconnecting strands. Based on their topological arrangements SBPs can divided into two classes (Fukami-Kobayashi et al., 1999) or more recently in seven clusters (Berntsson et al., 2010; Scheepers et al., 2016). Ligand binding takes place between the interface of the D1 and D2 subdomains and initiate a closure in a venus fly-trap mechanism (Mao and Mccammon, 1984), a key element in ligand recognition and signal transduction in all SBP associated protein families.

**Figure 1:**
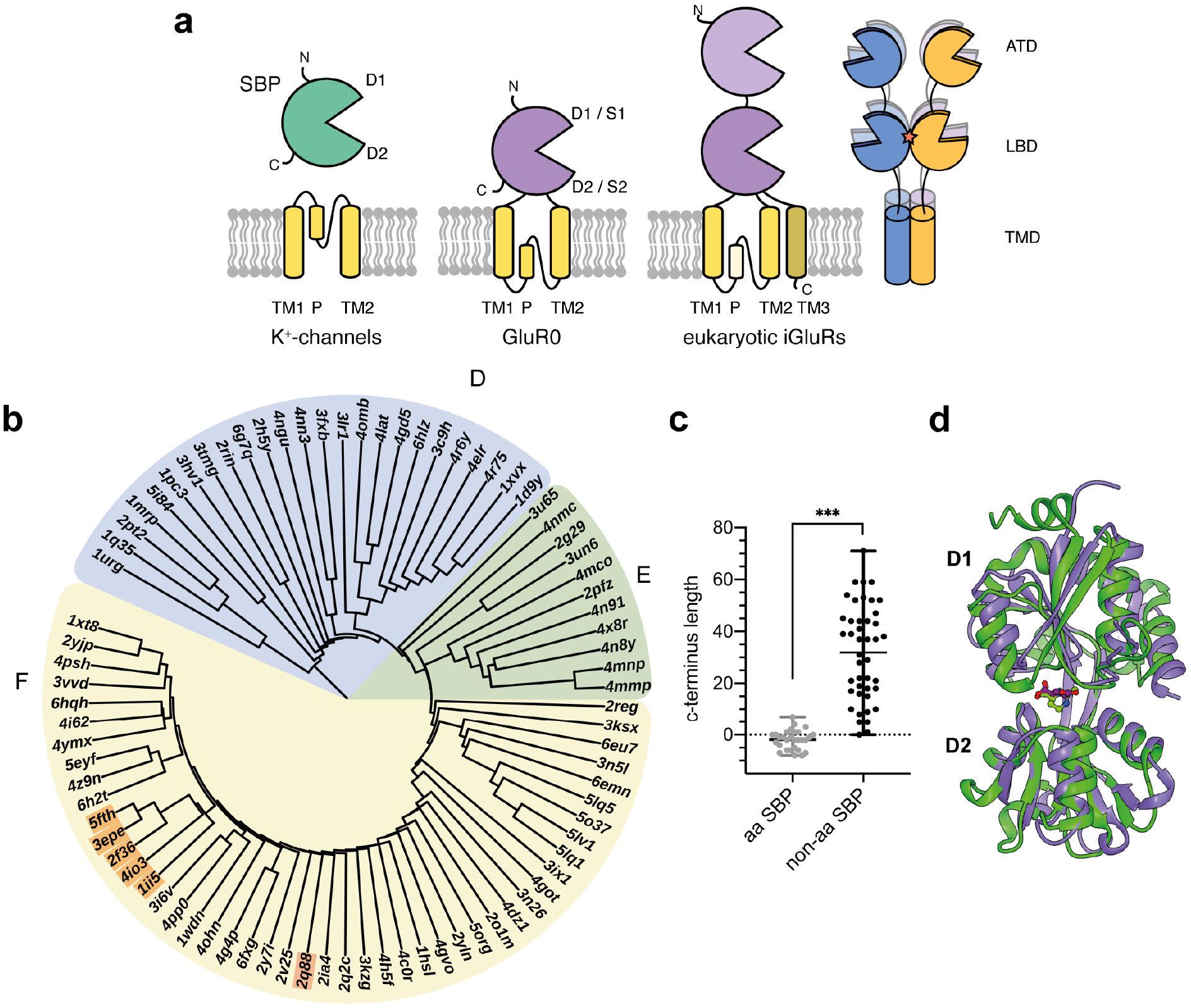
Structural relationship between iGluRs and SBP. **a** Modular organization of SBP, K^+^-channels and prokaryotic and eukaryotic iGluR (left) and tetrameric organization of metazoan iGluRs (right) including the amino-terminal domain (ATD), the ligand binding domain (LBD) with the dimerization interface (red star) and the transmembrane domain (TMD). **b** Structural distance tree of iGluR LBDs and SBPs structural homologous to the GluR0 LBD (1ii5). Clusters were categorized following the nomenclature of Berntsson et al. (Berntsson et al., 2010). The structural distance tree is subdivided into the following clusters: cluster F: two hinged SBPs including all iGluR LBDs (orange; GluR0:1ii5, AvGluR1:4io3, GluR5:2f36, GluR4:3epe, GluR2:5fth) and the ectoine binding protein EhuB (2q88), cluster E: SBDs from TRAP-transports cluster D: short hinged SBPs. **c** Comparison of the C-terminus lengths between amino acid (aa) or aa derivate SBPs and non-amino acid SBPs relatively measured to position Ser366 in the GluR0 LBD (p < 0.0001, Student’s two-tailed, unpaired t-test with Welch’s correction). **d** Structural superposition between the ligand-bound, closed conformations of the GluR0 LBD (purple, 1ii5) and the ectoine binding protein EhuB (green, 2q88).

Although iGluR LBDs and SBPs share some similarities in their overall structure and ligand recognition the functional compatibility of SBP to gate the channel pore of an iGluR has not been investigated. Here, we coupled in a proof of principle concept the bacterial ectoine binding protein EhuB to the channel pore of GluR0, building a functional ectoine activated receptor. With this approach, we provide the first experimental evidence that iGluRs originated by the fusion of an amino acid SBP to an ion channel. Further, our results highlight a conserved ligand binding mechanism in both protein families and the role of LBD dimerization in the functional evolution of iGluR from SBPs.

## Results

### Identification of an iGluR LBD structure-homologous SBP

The structural and molecular requirements for SBPs to build a LBD that is functionally linked with an (iGluR) channel pore to connect ligand binding with channel opening, which may provide insights into the evolution of iGluRs, remains widely unknown. To study these requirements, we wanted to create a functional chimeric receptor by replacing the LBD of the bacterial GluR0 receptor with a structurally related class II bacterial SBP. The bacterial GluR0 receptor was chosen due to its less complex architecture compared to eukaryotic iGluRs, characterized by the lack of an NTD, a M4 helix and an CTD, possibly displaying an evolutionary link between potassium channels, SBP and iGluRs (Chen et al., 1999). In the first step, we wanted to identify a SBP related to the GluR0 LBD. Due to the high number of SBPs and their low sequence identity, commonly below 20% (Berntsson et al., 2010), we decided to identify related bacterial SBPs based on their structural similarities using the Vast(+) algorithm (Madej et al., 2014). For protein identification, it must be considered that the extent of LBD closure must be conserved since it possibly contributes to a sufficient receptor activation (Neali Armstrong and Gouaux, 2000; Gill et al., 2008) and that additional SBDs that are part of enzyme complexes or gene regulators with different functional adaptions could be identified. Therefore, we used the structure of the glutamate-bound closed GluR0-LBD conformation (Mayer et al., 2001) (PDB ID: 1ii5) as a search template to ensure a similar extent of domain closure upon ligand binding (see Materials and Methods section). The 391 identified structures with root-mean-square deviation (RMSD) values up to 4 Å were further manually inspected to remove duplicates and open or unliganded structures, as well as structures of enzymes and gene regulators. To analyze the structural relationship and features between SBP and iGluR LBDs, the RMSD values of the 94 remaining SBPs and SBDs were further used to create a structural distance tree using the kitsch program (Fig. 1b, Supplementary Table S1). In general, the structures can be divided into 3 clusters according to Berntsson et al., 2010. The largest cluster F comprises of SBPs with two hinge segments (8-10 amino acids each) that bind a wide variety of substrates including amino acids, trigonal planar anions, compatible solutes and iron, and all iGluR LBDs. Interestingly, all iGluR LBDs form a small sub-cluster with a branching order supporting the current assumption with GluR0 representing a bacterial iGluR archetype and the freshwater bdelloid rotifer AvGluR1 (Janovjak et al., 2011) as a transitional stage between pro- and eukaryotic iGluRs. The third cluster D, which displays only SBPs, is characterized by two short hinge segments (4-5 amino acids each) and binds various substances like sugars, phosphates, iron and molybdate. Cluster E is placed between clusters D and F, exhibiting exclusively SBDs of TRAP- and TT-transporters. Interestingly, almost all SBPs that are structurally closely related to the iGluR bind amino acids or derivatives. Further analysis revealed that a common feature of this amino acid binding protein group is a short C-terminus with a similar length as found in GluR0, relatively to position Ser366 in the GluR0 LBD (Fig 1c), compared to non-amino acid binding proteins (p < 0.0001, Student’s two-tailed, unpaired t-test with Welch’s correction). The extended C-termini of non-amino acid binding proteins often attached to the upper D1-lobe, forming the LBD interface in iGluRs. Surprisingly, the ectoine binding protein EhuB is listed as the most related non-amino acid SBP to the GluR0 LBD within cluster II. EhuB is part of ABC transporter system Ehu from *Sinorhizobium meliloti*, that mediates the uptake of the compatible solutes ectoine and hydroxyectoine with high affinity (K_d_ = 1.6 μM) under osmotic stress (Hanekop et al., 2007). Despite a sequence identity of only 17 %, EhuB shows an almost identical overall structure to the GluR0 LBD with a RMSD of 2.2 Å and 216 of 258 superimposed amino acids (Figure 1c). Furthermore, both binding domains share a similar rotation angle in the closed, liganded conformation. These convenient features provided for us the basis of a proof-of-concept approach to combine the ectoine binding protein EhuB (PDB ID 2Q88) with the channel pore of GluR0. Since EhuB presumably evolved in absence of any adaptive constrains to gate the pore of ion channels it substantiate the general mechanistic and evolutional capability of SBPs to gate an ion channel by converting chemical information into an electric signal.

### Replacement of the GluR0 LBD with the ectoine binding protein

To examine whether GluR0 can serve as a foundation for functionally coupling the ectoine binding protein EhuB to its channel pore, we first expressed the codon optimized GluR0 receptor containing a GluR6 signal peptide and a N-terminal c-Myc tag in *Xenopus laevis* oocytes. Injected oocytes produced glutamate dependent inward currents at −80 mV in potassium ringer solution (100 mM KCl, 1 mM CaCl_2_, 5 mM HEPES, pH 7.4) with maximum peak currents of 162.5 ± 51.2 nA (mean ± SEM; n = 4) and an EC_50_ value of 29.9 ± 5 μM. After testing the applicability of GluR0 for receptor chimera design, we first asked if the fusion of the unmodified EhuB binding protein with the GluR0 TMD is sufficient to gate the channel pore. This chimeric receptor is further termed GluR0EhuB. In GluR0, the LBD is subdivided into two subdomains separated by the TMD and connected by two linkers. In the absence of a full-length GluR0 crystal structure, we estimated the connection position between the GluR0 linkers and the prospective EhuB LBD by superposition of the crystal structures of both binding domains (Fig. 2a,b). The previously described ends of the GluR0 LBD lobes (P139, A255) (Mayer et al., 2001) were set as a reference point and EhuB was fused at position K132 and N135 with the GluR0 S1-M1 and M2-S2 linkers. To maintain the original GluR0 linker length, positions K133, G134 in EhuB were deleted in the GluR0EhuB receptor. Additionally, the construct was codon optimized and the native EhuB signal peptide was replaced by a GluR6 signal peptide, followed by a c-Myc tag (see materials and methods section). The construct was expressed in oocytes and measured in potassium ringer solution without sodium chloride (100 mM KCl, 1 mM CaCl_2_, 5 mM HEPES, pH 7.4) at −80 mV. Unfortunately, we could not observe a ligand dependent current after the application of various ectoine concentrations (1 nM – 1 mM). Analyzing the total and surface expression by surface biotinylation and western blot (Fig. 2d) revealed a clear band at the calculated molecular weight of approximately 40 kDa. We therefore concluded that the GluR0EhuB chimera was expressed at the cell surface, indicating that receptor folding and assembly is not impaired.

**Figure 2:**
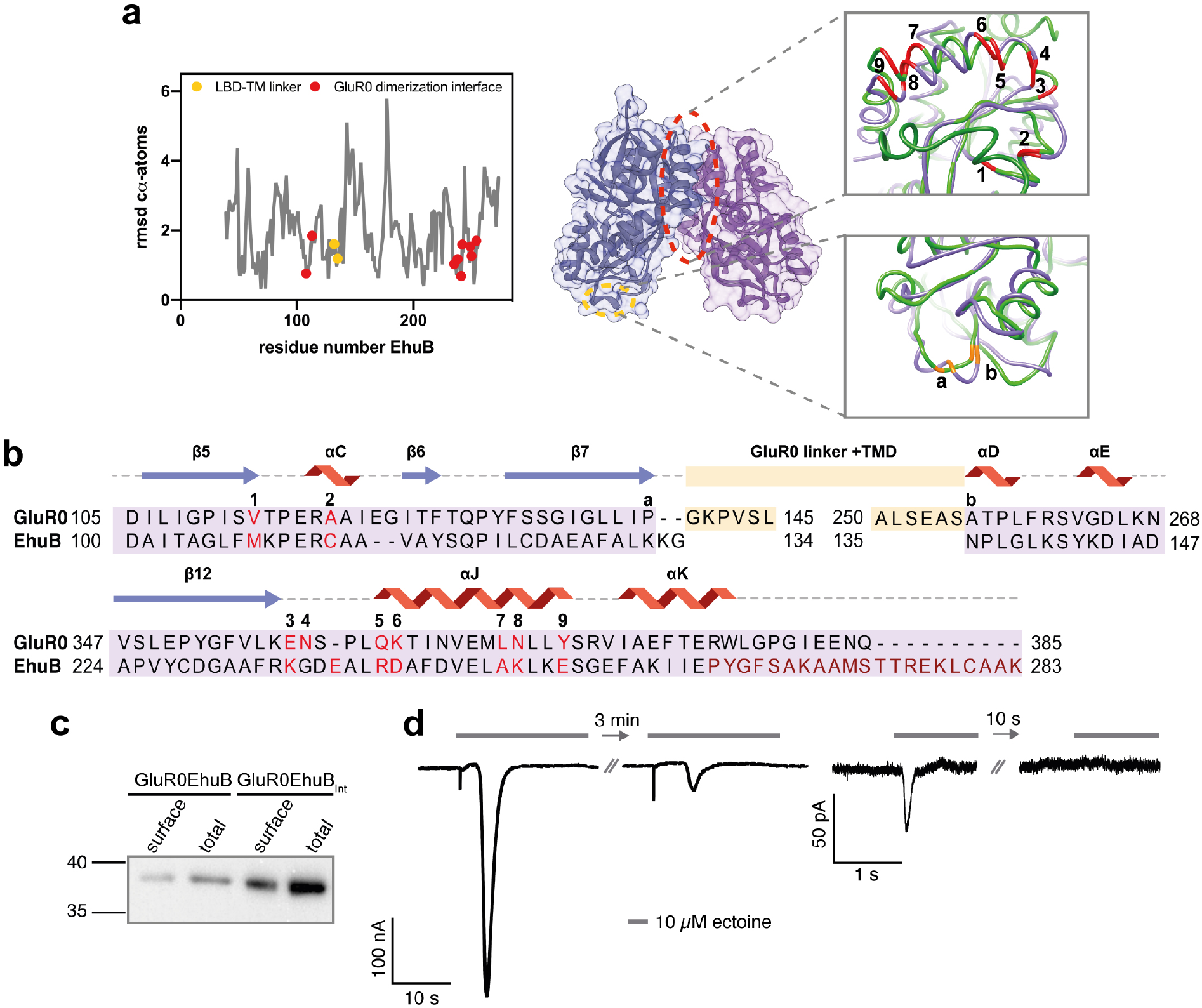
Design and expression of the chimeric GluR0EhuB and GluR0EhuBI_nt_ receptors. **a** Local RMSD of cα-atoms between the GluR0 LBD and EhuB (left) and structural overview (right) of the exchanged amino acid positions within the EhuB LBD in GluR0EhuB and GluR0EhuB_Int_. The connection points to the LBD-TMD linkers (orange) and amino acid positions that are responsible for LBD dimerization in GluR0 and are exchanged in GluR0EhuB_Int_ (red) are highlighted. **b** Structural sequence alignment of EhuB and the GluR0 LBD, including the linker positions (orange a,b), the substituted amino acid positions of the dimerization interface (red 1-9) and the deleted helices 10-12 (dark red) in GluR0EhuB_Int_. **c** Western blot of total and surface expression of GluR0EhuB (left) and GluR0EhuB_Int_ (right) with a calculated molecular weight of 40 kDa and 38 kDa, respectively. Proteins were detected using a c-Myc tag located at the N-terminus after the signal peptide. **d** Representative traces of whole-cell currents of GluR0EhuB_Int_ in *Xenopus* oocytes (left) and HEK293 cells (right) in response to 10 μM ectoine. Arrow is indicating the reduced receptor response to a second application of ectoine after 3 min or 10 s wash out.

### Optimization of the ectoine binding protein as LBD

The LBDs of iGluRs exhibit extensive intermolecular interactions within the D1 lobe, responsible for LBD dimerization between two adjacent subunits (N Armstrong and Gouaux, 2000; Sobolevsky et al., 2009c; Yelshanskaya et al., 2014). After ligand binding, the dimerization interface maintaining the D1-lobes in a back-to-back dimer arrangement, allowing the D2-lobes to separate from each other. This separation is a key step to induce the transduction of the conformational change within the LBD towards the membrane pore. Therefore, the LBD interface is crucial for channel activity by converting conformational changes within the LBD to channel opening, a feature that is probably missing in the bacterial EhuB SBP in the absence of any specific evolutionary adaption. In GluR0, the dimerization interface is characterized by several hydrogen bonds and van der Waals contacts mainly formed by 9 amino acids (V113, A118, E348, N349, Q353, K354, L361, N362, Y365) (Mayer et al., 2001). Further, EhuB exhibits an extended C-terminus, compared to the GluR0 LBD, formed by three helices (10-12), interacting with the same D1 area that promotes LBD dimerization in iGluRs. We speculated that the missing LBD dimer-interface and the large C-terminus in GluR0EhuB might prevent a specific D1-D1 LBD dimerization, resulting in a non-functional receptor chimera. Therefore, we decided to design a receptor chimera containing the GluR0 LBD dimer-interface in the fused ectoine-binding domain, hereinafter termed as GluR0EhuBInt (Int for interface). By structural superposition, we identified the amino acid positions in EhuB congruent with the amino acids mediating LBD dimerization in GluR0 (Fig. 2b,c). Due to the low deviations between the cα-atoms of both structures in the interface region (0.7 to 1.9 Å), we substituted the corresponding amino acids in the EhuB-binding domain (positions M108V, C113A, K235E, E238N, R241Q, D242K, A249L, K250N and E254Y according to EhuB) to reconstruct the GluR0 LBD dimerization interface. Since the C-terminal helices 10-12 in the original EhuB binding protein presumably interfere with LBD dimerization, we additionally deleted this part in the GluR0EhuB_Int_ receptor chimera. GluR0EhuB_Int_ was also well expressed and located at the cell surface, as indicated by western blot and surface biotinylation (Fig. 2d). Application of 10 μM ectoine (Fig. 2e), which lies in the physiological recognition range of EhuB, leaded to a ligand depended inward current of 347 ± 120 nA (mean ± SEM; n = 6) in oocytes and 79.3 ± 47.1 pA (mean ± SEM; n = 4) in HEK293 cells with a rapid inactivation. Unfortunately, receptor currents were tremendously reduced or not longer detectable after reapplying ectoine in a timespan of up to 20 min in oocytes or HEK293 cells. Our observations give evidence that the ectoine binding protein EhuB is capable to gate the channel pore of GluR0, leading to an ectoine dependent current with maximum currents comparable to GluR0. These findings imply that our chimeric receptor maintained its tetrameric assembly to form a functional ion channel that could be gated by the LBD upon ligand binding. Further, our data suggest that the formation of the inserted LBD dimerization interface is crucial to couple changes within the LBD to channel gating.

### Stabilization of the ectoine binding protein dimer interface

The insertion of a dimerization interface in GluR0EhuB_Int_ was sufficient to activate the receptor in the presence of ectoine. However, the modifications within the LBD in GluR0EhuB_Int_ resulted in a receptor with an impaired reactivation. We excluded the possibility that the ligand gets entrapped inside the binding domain after binding and cannot be washed out, caused by the introduced molecular changes in the ectoine-binding domain, since we did not observe a change in the leak current after receptor activation. Although the cα-backbones of the GluR0 LBD and EhuB are highly congruent, there are some minor structural differences in the exchanged amino acid positions, especially at C113A (1.9 Å), D242K (1.6 Å), A249L (1.6 Å) and E226Y (1.7 Å). It is likely that side chains of some substituted amino acids are not capable to form intermolecular interactions, weakening the dimerization interface. Therefore, we assumed that the dimerization interface gets disrupted to an extent incapable to reorganize after the first activation that traps the receptor in a closed, non-activable state. To test this hypothesis, we decided to strengthen the LBD dimerization interface by covalently coupling the D1-D1 subdomains by introducing a disulfide bond at position P119C and L376C (called GluR0EhuB_Int,P119C,L376C)_. This position (Fig. 3a) is functionally conserved in all AMPA- and Kainate-receptors and, if substituted with Cys or Tyr, prevents receptor desensitization by stabilizing the LBD dimers (L. Chen, K. Dürr, 2014; Sun et al., 2002; Weston et al., 2006). Application of ectoine (Fig. 3b, c) in GluR0EhuB_Int,P119C,L376C_ expressing HEK293 cells induced a concentration-dependent activation with a maximum current response of 65.1 ± 10.7 pA and an EC_50_ value of 14.1 ± 7.3 nM (mean ± SEM, n = 4). GluR0EhuB_Int,P119C,L376C_ evoked currents resembled some characteristics of GluR0, such as slow receptor activation, a high efficient dose-dependent activation, as well as a decreased desensitization upon prolonged applications (3 s) of ectoine, which is in agreement with previous studies on other members of the iGluR family (L. Chen, K. Dürr, 2014; Sun et al., 2002; Weston et al., 2006). Thus, by stabilizing the dimerization interface, we were able to retain receptor activity over multiple ectoine reapplications. Together, our results demonstrate the functional capability of the bacterial ectoine binding protein EhuB to gate the channel pore of GluR0 by specific changes in the amino acid sequence of the EhuB-binding domain, resulting in ligand and concentration dependent receptor currents.

**Figure 3:**
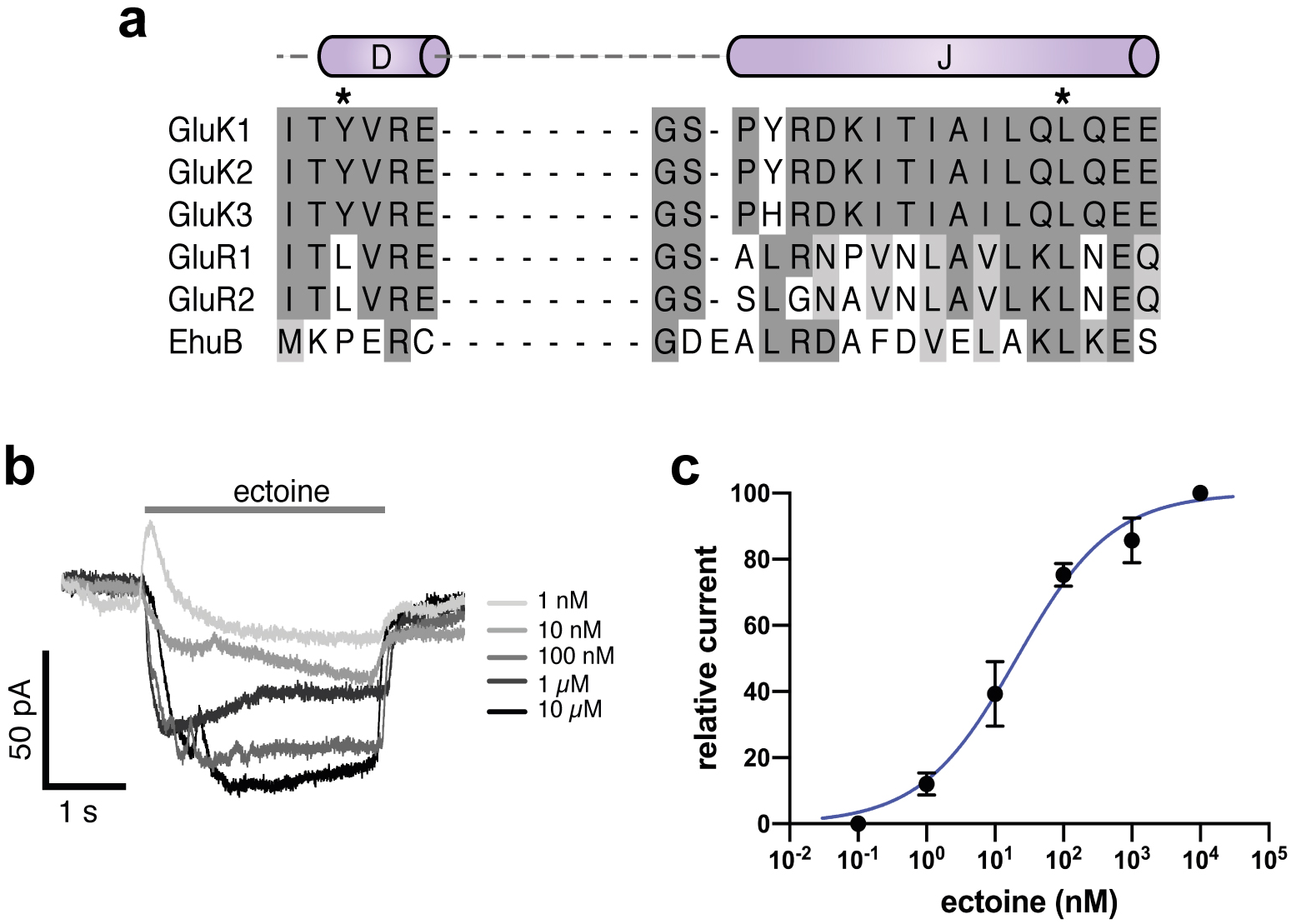
Stabilization of the LBD dimerization interface in GluR0EhuB_Int,P119C,L376C_ by disulfide bonds. **a** Sequence alignment of helices D and J of AMPA- and Kainate receptors that are involved in LBD dimerization interface formation. Previously published amino acid positions substituted with Cys or Tyr to covalently couple the D1 domain of each LBD-dimer in AMPA- and kainate recepors are highlighted with asterisks. **b** GluR0EhuB_Int,P119C,L376C_ responses to 3 s applications of 1 nM to 10 μM ectoine recorded from a single HEK293 cell. **c** Dose-response analysis of GluR0EhuB_Int,P119C,L376C_ revealed an EC_50_ value of 14.1 ± 7.3 nM (mean ± SEM, n = 4).

## Discussion

Although it has long been proposed that the LBDs of iGluRs are derived from bacterial SBPs (Felder et al., 1999), the evolutionary and functional compatibility of SBPs as an ancient module for ligand recognition in iGluRs has not yet been investigated. In this study, we characterized the basic molecular requirements to functionally couple a SBP to the channel pore of an iGluR, substantiating the functional compatibility and evolutionary kinship between both protein families. Our results indicate (i) a conserved ligand binding mechanism between SBPs and iGluR LBDs, (ii) that the formation of a LBD dimerization interface is a key step in iGluR evolution to couple ligand binding to channel gating and (iii) that iGluRs probably evolved by the fusion of class F amino acid binding proteins with potassium channels.

All iGluRs consist of at least two domains: the LBD and the ion channel-forming TMD. The common molecular architecture is most likely the result of different protein precursors, which fused during the evolution of iGluRs. It has been clearly shown that potassium channels display several similarities in their overall structure, topology and sequence with the pore domains of iGluRs (Chen et al., 1999; Kuner et al., 2003; Wood et al., 1995) and share a common gating mechanism (Schönrock et al., 2019), underscoring a common origin. Felder and colleagues first proposed over 20 years ago (Felder et al., 1999) that SBPs represent a blueprint of modern iGluR LBDs by sharing a common ligand binding mechanism. However, due to their high functional diversity and low sequence identity, it is insufficient to draw conclusions about a common origin of both protein domains. By coupling the ectoine binding protein EhuB to the channel pore of GluR0 in a semi-rational approach we emphasize a common origin and ligand binding mechanism of SBPs and iGluRs. In iGluRs, ligand binding can induce either a partial closure of the clamshell shaped LBD, as found in pre-active states, or a full closure that is associated with the active state. The extent of receptor activation depends on both the capability to induce a full LBD closure and the fraction of time the LBD occupies this fully closed conformation. In contrast, incomplete domain closure results in unproductive conformations and a decreased activity, as seen for partial agonists (Ahmed et al., 2011; Lau and Roux, 2011; Maltsev et al., 2008; Salazar et al., 2017; Twomey and Sobolevsky, 2018; Zhang et al., 2008). Interestingly, even SBPs sample a wide range of conformations of partial-closed structures, that are actively involved in substrate transport, as well as non-transported ligands adopting a distinct conformation to those that are actively transported (Boer et al., 2019; Gouridis et al., 2014; Tang et al., 2007). It is therefore evident that both the GluR0 LBD and EhuB must adopt a similar fully closured conformation in the presence of the TMD, to open the GluR0 channel pore. Furthermore, the functionality of our chimeric receptors suggests that the general mechanism of ligand binding and domain closure must be similar in EhuB and GluR0, exhibiting a comparable amount of free energy to stabilize the ligand and a similar probability to occupy the fully closed conformation. These findings highlight the general modular architecture of iGluRs and support the common hypothesis that the LBD originates from a bacterial SBP. It further suggests that the underlying mechanism of full- and partial agonism, as well as competitive inhibition in iGluRs, could be traced back to its bacterial precursors.

Further, our data sheds light on the minimum molecular requirements to obtain receptor function after a potential fusion event and govern the physiological properties of modern iGluR LBDs. The formation and maintenance of the back-to-back dimer arrangement by the D1-D1 interface is crucial for iGluR gating. During activation, the interface is responsible to adhere two adjacent LBDs. This allows the D2 lobes within the LBD dimers to separate from each other to transfer the conformational change via the linkers to the ion channel (Armstrong et al., 2006; Dürr et al., 2014; Sobolevsky et al., 2009a). In contrast, the same force that creates strain on the connecting linker upon ligand binding can lead to a partial or full rupture of the dimerization interface, known as desensitization. The rupture of the LBD dimerization interface results in a reorientation of the LBD dimer, allowing the D2 lobes to adopt a closed state-like conformation that disconnects the closed LBD conformation from channel activation (Meyerson et al., 2014; Plested, 2016; Schauder et al., 2013; Sun et al., 2002). Our data suggest that the bare fusion of a SBP to an ion channel pore is insufficient to form a functional receptor. We suppose that the TMD of the GluR0EhuB receptor assembles as a tetramer, since it is displayed at the plasma membrane, forming a putative functional channel pore. Since it has not been reported that EhuB assembles as dimers or oligomers in solution (Hanekop et al., 2007) and in the absence of any evolutionary pressure, it is unlikely that functional interactions are formed within the LBD layer of the GluR0EhuB receptor, holding the dimers in a fixed position during receptor gating. Therefore, we assume that ligand binding to the EhuB binding domain is decoupled from channel gating, since the force generated by clamshell closure cannot be used to separate the D2 lobes and transferred by the linkers. This is consistent with previous studies, showing that a complete rupture of the D1 dimerization interface by covalently connecting the D2 lobes, leads to a decoupling of agonist binding from ion channel gating (Armstrong et al., 2006). Remarkably, the change of a few amino acids within the putative LBD dimerization interface is sufficient to gain basic receptor activity, as seen in GluR0EhuB_Int_. However, the GluR0 LBD and EhuB protein are not completely homologues in their spatial structure. It is most likely that the inserted artificial dimerization interface is too weak to prevent a full rupture of the interface upon ligand binding that locks the receptor in a non-active conformation where ligand binding is disengaged from channel gating. This is in agreement that even minor perturbations of the dimerization interface by single point mutations enhance desensitization and can lead to a delayed receptor recovery (Fleck et al., 2003; Horning and Mayer, 2004; Partin et al., 1996; Sun et al., 2002). By covalently stabilizing the LBD dimerization interface in GluR0EhuB_Int,P119C,L376C_, we demonstrated that the impaired activation mechanism can be overcome and resulted in a receptor chimaera that is capable to undergo a full gating cycle from activation to deactivation and recovery. Moreover, the activation and deactivation kinetics observed in GluR0EhuB_Int,P119C,L376C_ resemble the slow activation of GluR0 and the constitutive activity upon a prolonged ligand application observed in LBD stabilized AMPA- and kainite receptors. In conclusion, these findings reveal the dimerization interface as the key element in the molecular evolution of iGluRs that is crucial to couple the clamshell closure of the binding domain to ion channel gating. The requirement of only minor adaptions to form a LBD dimerization interface to obtain a functional iGluR archetype is presumably an advantage in the functional evolution and physiological adaptation of iGluRs to fulfill different tasks among bacteria, plants, amoebas and metazoans.

The structural distance tree only provides limited information about the direct phylogenetic relationship of proteins, since it is only based on structural derivations and not on sequence comparisons. However, structural derivations are a result of changes in their protein sequence. We therefore can make some assumptions about the evolutionary origin of iGluR LBDs. It should be emphasized that the closest structural related SBPs in cluster F all bind amino acids or amino acid derivatives. This finding is less surprising since GluR0 as well as AvGluR1, two putative links between the evolution of modern metazoan iGluRs from SBPs and potassium channels, are activated by several polar amino acids (Chen et al., 1999; Janovjak et al., 2011). Furthermore, the LBD of AMPA receptors and GluR0 share a weak amino acid sequence homology with the glutamine binding protein from *Escherichia coli* (Chen et al., 1999). For that reason, it is most likely that the ancient SBP precursor belongs to class F SBPs. It is notable that all identified class F amino acid and amino acid derivate SBPs display a relatively short C-terminus compared to all non-amino acid binding proteins found in the more distant classes D and E. These extended C-termini often pack to the D1 lobe, masking the same helices that mediate LBD dimerization in iGluRs. Presumably, it was advantageous in the evolution of iGluRs, that the putative LBD dimer interface was freely accessible in amino acid binding proteins that facilitated a rapid functional adaption of dimer-dimer interactions without extensive changes in the protein structure. Moreover, the short C-termini of amino acid binding proteins possibly served as a putative S2-M4 linker to couple the additional M4 helix in later evolutionary iGluR stages.

In summary, by functionally coupling the ectoine binding protein EhuB to the GluR0 channel pore we substantiate the compatibility between both protein families and shedding light on the functional and molecular evolution of iGluRs from bacterial SBPs and potassium channels. Our results may help in the mechanistic understanding of ligand recognition in both protein families and drug design. Finally, our approach, combined with the diversity of recognized ligands and the structural adaptability of SBPs could be used as a versatile tool in the design of biosensor with a high specificity and efficiency.

## Materials and Methods

### Identification of homologous SBP and structural distance tree

For structure identification and building a structural distance tree, we used the previously described method from Scheepers et al., 2016 with some modifications. To identify SBPs that are structurally homologous to the LBD of GluR0, we used the ligand-bound LBD conformation (1ii5) as a search template against the Molecular Modeling Database (MMDB) using the Vector Alignment Search Tool (VAST+) (Madej et al., 2014). The resulting 354 PDB codes were matched against the UniProt IDs to filter out multiple structures of the same protein. Additionally, unliganded structures, enzymes and gene regulators were filtered out by manual inspection. This search resulted in about 94 SBP- and iGluR LBD-structures similar to the GluR0 LBD (Supplementary Table S1). The remaining structures were pairwise aligned using the PDBeFold server (Krissinel and Henrick, 2004). To build the structural distance tree, the RMSD values obtained from PBDeFold were loaded into the KITSCH program of the PHYLIPS package with default parameters (Felsenstein, 2004). Protein clusters were verified by visual inspection using UCFS Chimera (Pettersen et al., 2004) according to Berntsson et al., 2010 and the structural distance tree was visualized using iTOL (https://itol.embl.de/). For the determining the C-termini lengths of amino acid and non-amino acid SBP, structures were pairwise aligned to the GluR0 LBD (1ii5) and inspected using UCSF Chimera. The relative C-terminus was specified relative to position Ser366 in the GluR0 LBD. Statistical significance was determined at the p ≤ 0.05 (*), p ≤ 0.01 (**), p ≤ 0.001 (***) and p ≤ 0.01 (****), levels using a Student’s two-tailed, unpaired t-test with Welch’s correction. Equality of variances was tested using a F-test.

### Protein engineering

In this study, we used the sequences from GluR0 (UniProt ID P73797) and EhuB (UniProt ID Q92WC8). Signal peptide cleavage site was predicted using SignalP v. 4.1 (Nielsen et al., 1997). The intrinsic GluR0 (residues 1-19) or EhuB signal peptides (residues 1 – 27) were replaced by the GluR6 signal peptide (MRIICRQIVLLFSGFWGLAMG), followed by a c-myc tag (EQKLISEED) and a short linker (SGTPT). For GluR0EhuB, EhuB D1 domain (residues 28 – 132) was fused to the N-terminal and the D2 domain (residues 135 – 283) to the C-terminal end of the GluR0 TMD, including the linker (residues 139 – 255). For GluR0EhuB_Int_, the positions M81V, C86A, K208E, E211N, R214Q, D215K, A222L, K223N and E226Y were substituted and residues 266 – 283 were deleted (positions corresponding to the original EhuB protein). For receptor expression in eukaryotic expression systems, the sequences of all constructs were codon-optimized *for Xenopus laevis.* GluR0, GluR0EhuB and GluR0EhuB_Int_ DNA strings were synthesized (GeneArt, Thermo Fisher Scientific, Regensburg, Germany) and subsequently cloned into the vector pCDNA3.1(+) via included *NheI* and *XhoI* restriction sites. For GluR0EhuB_Int,P119C,L376C_, the primers P119C forward (ATTGTTTGTCAAGTGTGAGAGAGCCGCT) and reverse (AGCGGCTCTCTCACACTTGACAAACAAT), as well as L376C forward (TGATGTTGAACTCTGTAATCTGAAATACTCCG) and reverse (CGGAGTATTTCAGATTACAGAGTTCAACATCA) were used for mutagenesis polymerase chain reaction (PCR). PCR reaction parameters were as follows: initial denaturation at 95 °C for 60 s; 30 cycles at 95 °C and 30 s, 56 °C for 15 s and 72 °C for 150 s; and one cycle at 72 °C for 180 s. All constructs were confirmed by sequencing (Seqlab, Göttingen, Germany).

### Sequences

GluR0EhuB:

MRIICRQIVLLFSGFWGLAMGEQKLISEEDLSGTPTDENKLEELKEQGFARIAIANEPPFTAVGADGKVSGAAPDVAREIFKRLGVADVVASISEYGAMIPGLQAGRHDAITAGLFMKPERCAAVAYSQPILCDAEAFALKGKPVSLWERFSPFFGIAALSSAGVLTLLLFLVGNLIWLAEHRKNPEQFSPHYPEGVQNGMWFALVTLTTVGYGDRSPRTKLGQLVAGVWMLVALLSFSSITAGLASAFSTALSEASNPLGLKSYKDIADNPDAKIGAPGGGTEEKLALEAGVPRDRVIVVPDGQSGLKMLQDGRIDVYSLPVLSINDLVSKANDPNVEVLA PVEGAPVYCDGAAFRKGDEALRDAFDVELAKLKESGEFAKIIEPYGFSAKAAMSTTREKLCAAK

GluR0EhuB_Int_

MRIICRQIVLLFSGFWGLAMGEQKLISEEDLSGTPTDENKLEELKEQGFARIAIANEPPFTAVGADGKVSGAAPDVAREIFKRLGVADVVASISEYGAMIPGLQAGRHDAITAGLFVKPERAAAVAYSQPILCDAEAFALKGKPVSLWERFSPFFGIAALSSAGVLTLLLFLVGNLIWLAEHRKNPEQFSPHYPEGVQNGMWFALVTLTTVGYGDRSPRTKLGQLVAG VWMLVALLSFSSITAGLASAFSTALSEASNPLGLKSYKDIADNPDAKIGAPGGGTEEKLALEAGVPRDRVIVVPDGQSGLKMLQDGRIDVYSLPVLSINDLVSKANDPNVEVLA PVEGAPVYCDGAAFREGDNALQKAFDVELLNLKYSGEFAKIIEPYG

### Heterologous expression in X. laevis oocytes

Constructs were expressed in *X. laevis* oocytes as described previously (Schönrock et al., 2019). In brief, all constructs in pCDNA3.1(+) were linearized with NotI and cRNA was synthesized using the AmpliCap-Max™ T7 High Yield Message Maker Kit (Cellscript, Madison, WI, USA). Surgically obtained oocytes from female *X. laevis* were enzymatically separated and defoliculated using 0.8 mg collagenase in ringer solution (96mM NaCl, 2 mM KCl, 1mM CaCl_2_, 1mM MgCl2, 5mM HEPES; pH 7.4) for 12 h and reaction was stopped with Ca^2+^-free ringer solution. 50 ng of capped and polyadenylated RNA with 50 nl water was microinjected in defoliculated stage IV – V oocytes and incubated 3 – 7 days in ND-96 solution (96 mM NaCl, 2 mM KCl, 1 mM MgCl2, 5 mM HEPES, 50-mg ml-1 gentamycin; pH 7.5) at 18 °C. Related oocyte experiments (for example total and surface expression) were performed with the same oocyte batches on the same day.

### Heterologous expression in HEK293 cells

HEK293 cells were cultured in minimum essential medium (MEM) supplemented with 10 % (v/v) FCS, 2 mM L-glutamine and streptomycin (100 μg/ml). Before transfection, 5×10^5^ cells were reseeded into T25 flask and transfected using TurboFect (Thermo Fisher Scientific, Waltham, MA, USA) and 4 μg plasmid DNA per flask of the respective construct in pCDNA3.1(+) and pEGFP-N1 as a transfection control with a ratio of 4:1. After transfection, cells were incubated for 48-72 h at 37 °C and 5% CO_2_.

### Surface biotinylation

For surface biotinylation, 20 oocytes per construct were injected as described above and incubated for 3 days in ND-96 solution at 18 °C. Oocytes were washed three times with PBS (100 mM phosphate buffer, 150 mM NaCl; pH 7.2) and surface proteins were biotinylated using 0.5 mg/ml Sulfo-NHS-SS-Biotin (Thermo Fisher Scientific) in PBS for 40 min at room temperature. Reaction was stopped with 50 mM Tris/HCl; pH 8.0. Oocytes were mechanically homogenized and membrane proteins were extracted using DDM (100 mM phosphate buffer, 150 mM NaCl 0.5 % n-dodecyl-β-D-maltoside, SIGMAFAST Protease Inhibitor Cocktail (Sigma-Aldrich, St. Louis, MO, USA); pH 8.0) for 15 min at 4°C. Biotinylated membrane proteins were purified using Streptavidin High Performance Spintrap columns (GE Healthcare, Chicago, IL, USA).

### Western blotting

For total protein expression, 10 oocytes per construct were injected and incubated for 3 days. Oocytes were washed with 0.1 M sodium phosphate buffer, pH 8.0 and mechanically homogenized. Membrane proteins were solubilized with 25 μl lysis buffer (0.1 M sodium phosphate buffer pH 8.0, 0.5 % DDM, 0.01% pefa block) for 10 min at 4 °C and subsequently centrifuged with 1000 x g at 4°C. HEK293 cells were transfected by electroporation as described above and incubated 48 h. Cells were scraped, lysed with 200 μl lysis buffer and subsequently centrifuged 20 min with 16000 xg at 4°C. For SDS-PAGE and western blot, total and surface labeled proteins were separated using a 10% sodium dodecyl sulfate polyacrylamide gel electrophoresis (SDS-PAGE) and transferred to a PVDF membrane (Bio-Rad, Feldkirchen, Germany). The membrane was blocked for 1 h in PBS-T supplemented with 5% skim milk and afterward incubated with 1:600 primary c-Myc-tag Polyclonal antibody (sc-789, Santa Cruz Biotechnology, Dallas, TX, USA) in PBS-T containing 1% skim milk over night at 4 °C. After washing with PBS-T, the secondary goat anti-rabbit IgG-HRP (1:10000) (sc-2054, Santa Cruz Biotechnology) was incubated in TBS-T containing 1% skim milk for 1 h at room temperature. The membrane was washed three times in TBS-T and the signal was visualized using Pierce Western Blotting Substrate (Thermo Fisher Scientific) and detected with a CCD camera.

### Electrophysiological recordings of X. laevis oocytes using two-electrode voltage clamp

3-7 days after injection, whole-cell currents were recorded by two-electrode voltage-clamp at −80 mV using microelectrodes filled with 3M KCl (resistance 0.8–2.5 MΩ). Data were sampled at 5 kHz after low-pass filtering at 200 Hz using an Axoclamp 900A amplifier connected to a Digidata 1550A digitizer and recorded with Clampex 10.7 (Molecular Devices, San Jose, USA). Oocytes were placed in a perfusion chamber and rinsed with high potassium Ringer’s solution (100 mM KCl, 1 mM CaCl_2_, 5 mM HEPES, pH 7.4 with KOH). Ectoine (10 μM to 0.1 nM) or L-glutamate (1 mM to 1 μM) was applied to the oocytes in external solution. For dose-response analysis, normalized current responses were plotted against the agonist concentration and fitted with a sigmoidal Hill equation *I*/*I*_*max*_ = 100 × *x*^*h*^ + *EC*_50_^*h*^ in GraphPad Prism 9 (GraphPad Software Inc., La Jolla, USA) where I/I_max_ is the normalized current, x the concentration, h the Hill coefficient and EC_50_ the agonist concentration resulting in a half-maximal response. Data and graphs presented in mean ± SEM.

### Electrophysiological recordings using Port-a-patch

Electrophysiological recordings in HEK293 cells were performed using the Port-a-Patch system (Nanion Technologies, München, Germany). Adherent HEK293 cells were harvested 48-72 h after transfection using 1% accutase solution. The reaction was stopped with MEM and the cell suspension was centrifuged at 100 xg for 3 min. Cells were resuspended in extracellular solution (140 mM KCl, 1 mM MgCl_2_, 2 mM CaCl_2_, 5 mM D-glucose, 10 mM HEPES; pH 7.4; osmolarity 298 mOsmol) with a final concentration of 2×10^6^ - 3×10^6^ cells/ml. Measurements were performed using a Port-a-Patch system with external perfusion system (Nanion Technologies) and NPC-1 chips (3-5 MΩ) according to the manufacturers instructions, connected to an EPC10 amplifier (HEKA, Ludwigshafen, Germany), PatchControl (Nanion Technologies) and Patchmaster software (HEKA). External solution and internal solution (50 mM KCl, 10 mM NaCl, 60 mM KF, 20 mM EGTA, 1 mM ATP, 10 mM HEPES; pH 7.2; osmolarity 285 mOsmol) were used for experiments and seal enhancer (80 mM NaCl, 3 mM KCl, 10 mM MgCl2, 35 mM CaCl_2_, 10 mM HEPES; pH 7.4; osmolarity 298 mOsmol) was used for sealing cells. For recording receptor currents, cells were clamped at −60 mV and continuously perfused with external solution during recording. Ectoine dilution series (10 μM – 0.1 nM) was dissolved in external solution. Agonist was applied for 3 s, followed by 10 s of wash. Dose-response analysis was performed as described above. Data and graphs presented in mean ± SEM.

## Acknowledgements

This work has been supported in the frame of the LOEWE project iNAPO by the Hessen State Ministry of Higher Education, Research and the Arts. The authors acknowledge support by the German Research Foundation and the Open Access Publishing Fund of Technische Universität Darmstadt.

## Abbreviations

AMPA: α-amino-3-hydroxy-5-methyl-4-isoaxazolepropionic acid
CTD: C-terminal domain
iGluR: ionotropic glutamate receptor
LBD: ligand binding domain
NTD: N-terminal domain
RMSD: root-mean-square deviation
SBD: substrate binding domain
SBP: substrate binding protein
TMD: transmembrane domain

---

**Supplementary Table S1.**
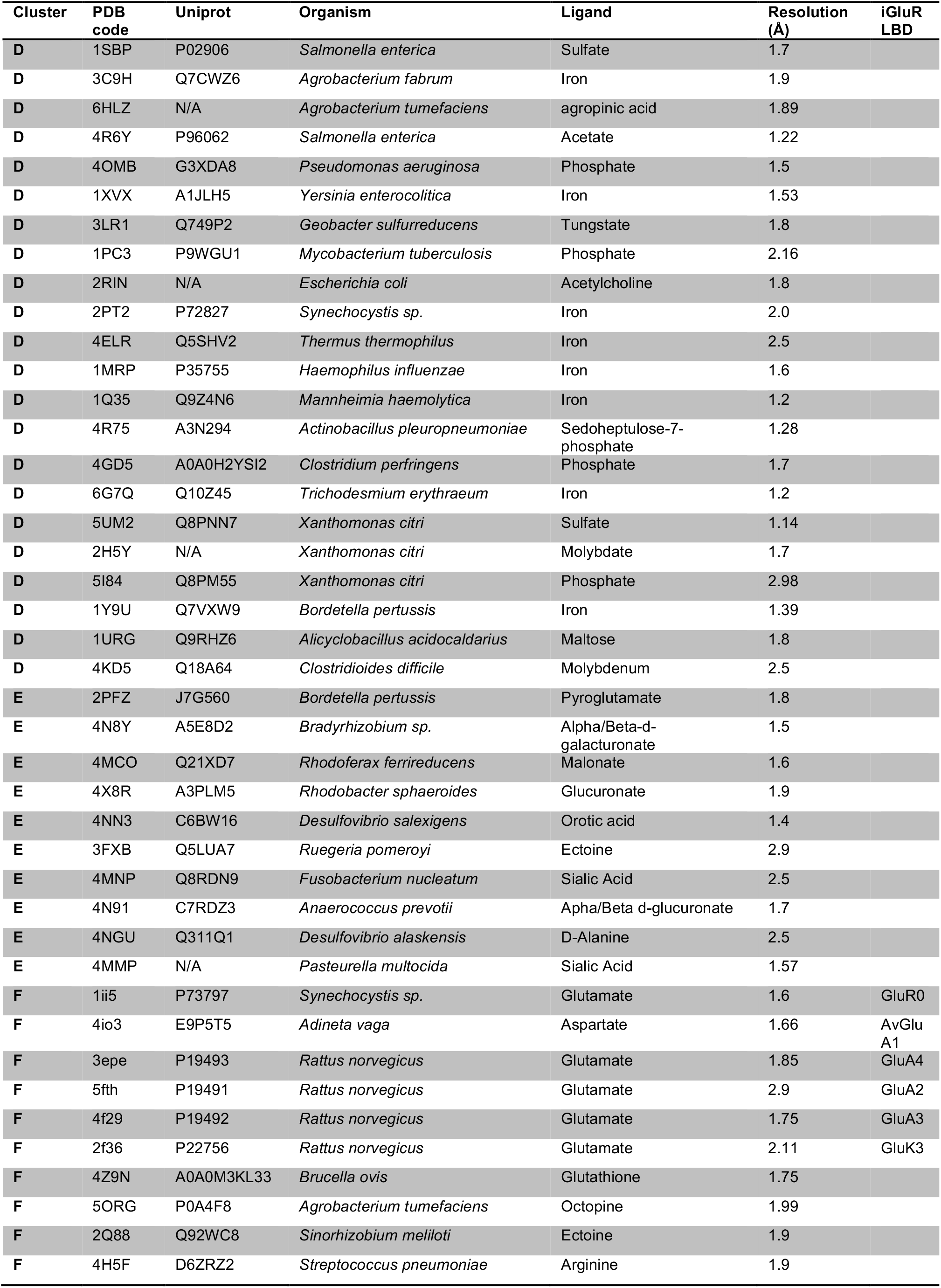
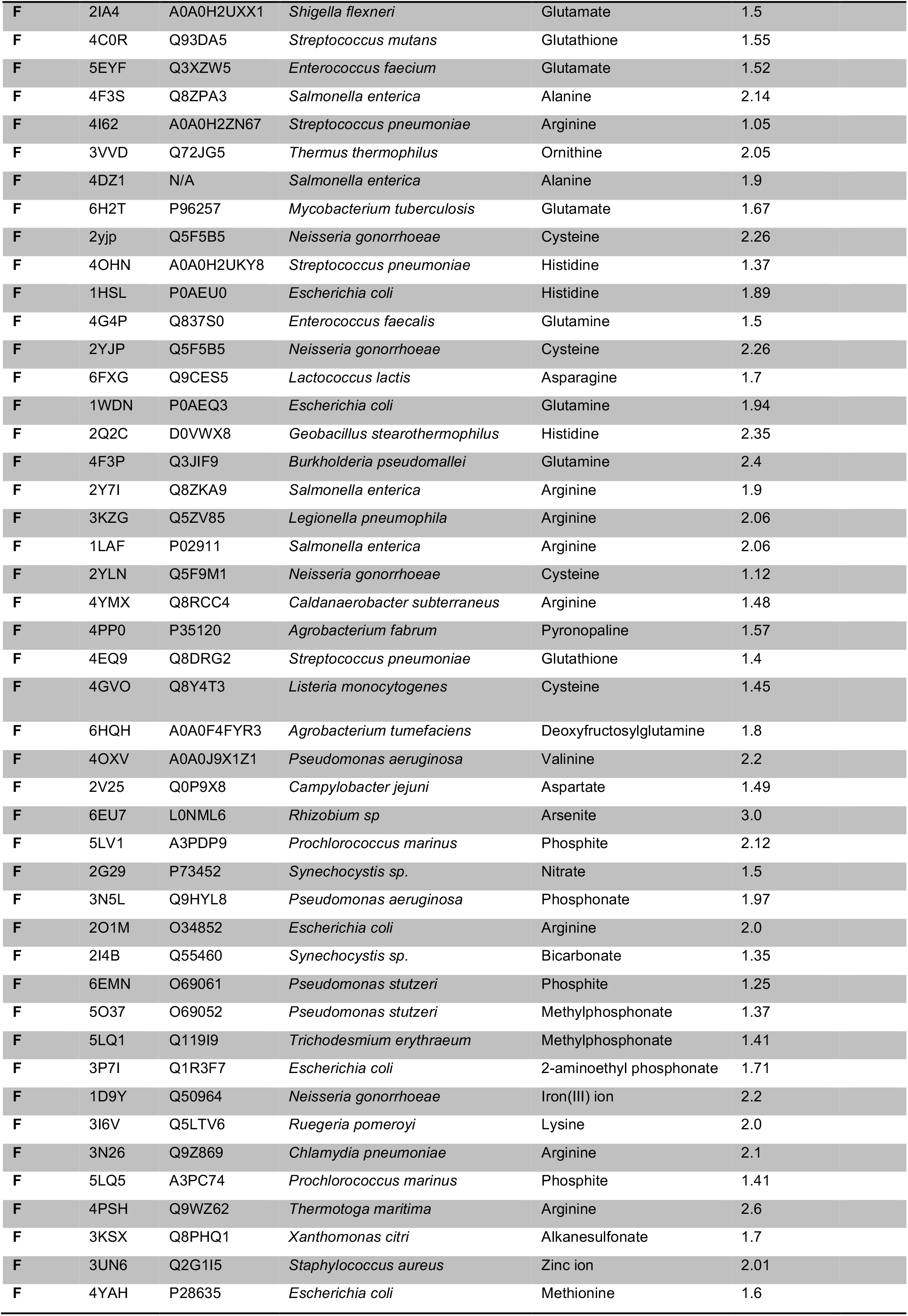
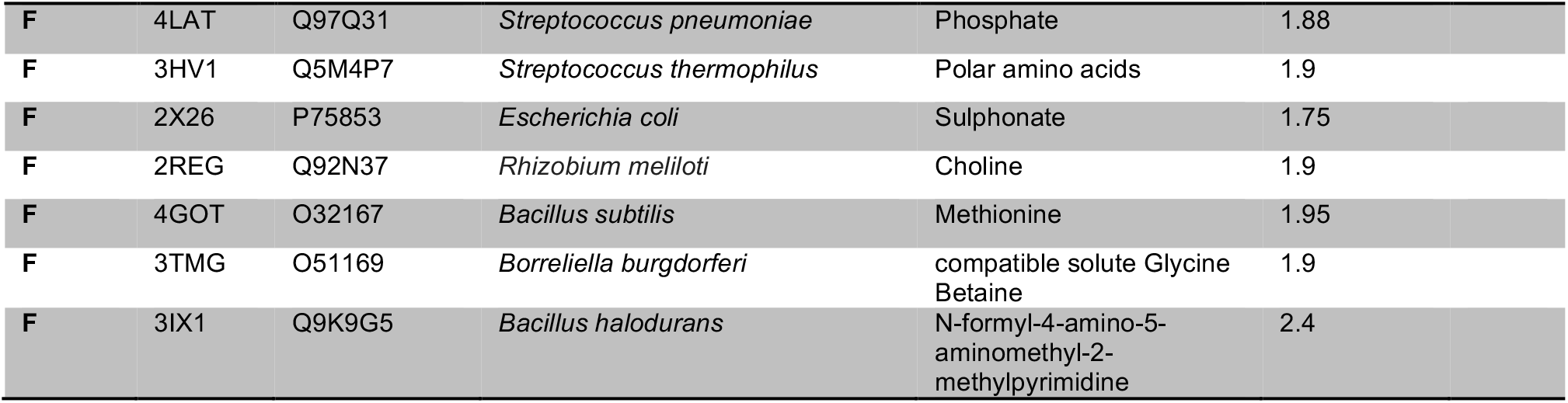
Overview of GluR0 (1ii5) structure homologoussubstrate-binding proteins identified in the Molecular Modeling Database (MMDB) using the Vector Alignment Search Tool (VAST+) and used fort he structural distance tree.

| Cluster | PDB code | Uniprot | Organism | Ligand | Resolution (Å) | iGluR LBD |
| --- | --- | --- | --- | --- | --- | --- |
| D | 1SBP | P02906 | *Salmonella enterica* | Sulfate | 1.7 |  |
| D | 3C9H | Q7CWZ6 | *Agrobacterium fabrum* | Iron | 1.9 |  |
| D | 6HLZ | N/A | *Agrobacterium tumefaciens* | agropinic acid | 1.89 |  |
| D | 4R6Y | P96062 | *Salmonella enterica* | Acetate | 1.22 |  |
| D | 4OMB | G3XDA8 | *Pseudomonas aeruginosa* | Phosphate | 1.5 |  |
| D | 1XVX | A1JLH5 | *Yersinia enterocolitica* | Iron | 1.53 |  |
| D | 3LR1 | Q749P2 | *Geobacter sulfurreducens* | Tungstate | 1.8 |  |
| D | 1PC3 | P9WGU1 | *Mycobacterium tuberculosis* | Phosphate | 2.16 |  |
| D | 2RIN | N/A | *Escherichia coli* | Acetylcholine | 1.8 |  |
| D | 2PT2 | P72827 | *Synechocystis* sp. | Iron | 2.0 |  |
| D | 4ELR | Q5SHV2 | *Thermus thermophilus* | Iron | 2.5 |  |
| D | 1MRP | P35755 | *Haemophilus influenzae* | Iron | 1.6 |  |
| D | 1Q35 | Q9Z4N6 | *Mannheimia haemolytica* | Iron | 1.2 |  |
| D | 4R75 | A3N294 | *Actinobacillus pleuropneumoniae* | Sedoheptulose-7-phosphate | 1.28 |  |
| D | 4GD5 | A0A0H2YSI2 | *Clostridium perfringens* | Phosphate | 1.7 |  |
| D | 6G7Q | Q10Z45 | *Trichodesmium erythraeum* | Iron | 1.2 |  |
| D | 5UM2 | Q8PNN7 | *Xanthomonas citri* | Sulfate | 1.14 |  |
| D | 2H5Y | N/A | *Xanthomonas citri* | Molybdate | 1.7 |  |
| D | 5I84 | Q8PM55 | *Xanthomonas citri* | Phosphate | 2.98 |  |
| D | 1Y9U | Q7VXW9 | *Bordetella pertussis* | Iron | 1.39 |  |
| D | 1URG | Q9RHZ6 | *Alicyclobacillus acidocaldarius* | Maltose | 1.8 |  |
| D | 4KD5 | Q18A64 | *Clostridioides difficile* | Molybdenum | 2.5 |  |
| E | 2PFZ | J7G560 | *Bordetella pertussis* | Pyroglutamate | 1.8 |  |
| E | 4N8Y | A5E8D2 | *Bradyrhizobium* sp. | Alpha/Beta-d-galacturonate | 1.5 |  |
| E | 4MCO | Q21XD7 | *Rhodoferrax ferrireducens* | Malonate | 1.6 |  |
| E | 4X8R | A3PLM5 | *Rhodobacter sphaeroides* | Glucuronate | 1.9 |  |
| E | 4NN3 | C6BW16 | *Desulfovibrio salexigens* | Orotic acid | 1.4 |  |
| E | 3FXB | Q5LUA7 | *Ruegeria pomeroyi* | Ectoine | 2.9 |  |
| E | 4MNP | Q8RDN9 | *Fusobacterium nucleatum* | Sialic Acid | 2.5 |  |
| E | 4N91 | C7RDZ3 | *Anaerococcus prevotii* | Apha/Beta d-glucuronate | 1.7 |  |
| E | 4NGU | Q311Q1 | *Desulfovibrio alaskensis* | D-Alanine | 2.5 |  |
| E | 4MMP | N/A | *Pasteurella multocida* | Sialic Acid | 1.57 |  |
| F | 1ii5 | P73797 | *Synechocystis* sp. | Glutamate | 1.6 | GluR0 |
| F | 4io3 | E9P5T5 | *Adineta vaga* | Aspartate | 1.66 | AvGlu A1 |
| F | 3epe | P19493 | *Rattus norvegicus* | Glutamate | 1.85 | GluA4 |
| F | 5fth | P19491 | *Rattus norvegicus* | Glutamate | 2.9 | GluA2 |
| F | 4f29 | P19492 | *Rattus norvegicus* | Glutamate | 1.75 | GluA3 |
| F | 2f36 | P22756 | *Rattus norvegicus* | Glutamate | 2.11 | GluK3 |
| F | 4Z9N | A0A0M3KL33 | *Brucella ovis* | Glutathione | 1.75 |  |
| F | 5ORG | P0A4F8 | *Agrobacterium tumefaciens* | Octopine | 1.99 |  |
| F | 2Q88 | Q92WC8 | *Sinorhizobium meliloti* | Ectoine | 1.9 |  |
| F | 4H5F | D6ZRZ2 | *Streptococcus pneumoniae* | Arginine | 1.9 |  |

|  |  |  |  |  |  |
| --- | --- | --- | --- | --- | --- |
| F | 2IA4 | A0A0H2UXX1 | *Shigella flexneri* | Glutamate | 1.5 |
| F | 4C0R | Q93DA5 | *Streptococcus mutans* | Glutathione | 1.55 |
| F | 5EYF | Q3XZW5 | *Enterococcus faecium* | Glutamate | 1.52 |
| F | 4F3S | Q8ZPA3 | *Salmonella enterica* | Alanine | 2.14 |
| F | 4I62 | A0A0H2ZN67 | *Streptococcus pneumoniae* | Arginine | 1.05 |
| F | 3VVD | Q72JG5 | *Thermus thermophilus* | Ornithine | 2.05 |
| F | 4DZ1 | N/A | *Salmonella enterica* | Alanine | 1.9 |
| F | 6H2T | P96257 | *Mycobacterium tuberculosis* | Glutamate | 1.67 |
| F | 2yjp | Q5F5B5 | *Neisseria gonorrhoeae* | Cysteine | 2.26 |
| F | 4OHN | A0A0H2UKY8 | *Streptococcus pneumoniae* | Histidine | 1.37 |
| F | 1HSL | P0AEU0 | *Escherichia coli* | Histidine | 1.89 |
| F | 4G4P | Q837S0 | *Enterococcus faecalis* | Glutamine | 1.5 |
| F | 2YJP | Q5F5B5 | *Neisseria gonorrhoeae* | Cysteine | 2.26 |
| F | 6FXG | Q9CES5 | *Lactococcus lactis* | Asparagine | 1.7 |
| F | 1WDN | P0AEQ3 | *Escherichia coli* | Glutamine | 1.94 |
| F | 2Q2C | D0VWX8 | *Geobacillus stearothermophilus* | Histidine | 2.35 |
| F | 4F3P | Q3JIF9 | *Burkholderia pseudomallei* | Glutamine | 2.4 |
| F | 2Y7I | Q8ZKA9 | *Salmonella enterica* | Arginine | 1.9 |
| F | 3KZG | Q5ZV85 | *Legionella pneumophila* | Arginine | 2.06 |
| F | 1LAF | P02911 | *Salmonella enterica* | Arginine | 2.06 |
| F | 2YLN | Q5F9M1 | *Neisseria gonorrhoeae* | Cysteine | 1.12 |
| F | 4YMX | Q8RCC4 | *Caldanaerobacter subterraneus* | Arginine | 1.48 |
| F | 4PP0 | P35120 | *Agrobacterium fabrum* | Pyronopaline | 1.57 |
| F | 4EQ9 | Q8DRG2 | *Streptococcus pneumoniae* | Glutathione | 1.4 |
| F | 4GVO | Q8Y4T3 | *Listeria monocytogenes* | Cysteine | 1.45 |
| F | 6HQH | A0A0F4FYR3 | *Agrobacterium tumefaciens* | Deoxyfructosylglutamine | 1.8 |
| F | 4OXV | A0A0J9X1Z1 | *Pseudomonas aeruginosa* | Valinine | 2.2 |
| F | 2V25 | Q0P9X8 | *Campylobacter jejuni* | Aspartate | 1.49 |
| F | 6EU7 | L0NML6 | *Rhizobium sp* | Arsenite | 3.0 |
| F | 5LV1 | A3PDP9 | *Prochlorococcus marinus* | Phosphite | 2.12 |
| F | 2G29 | P73452 | *Synechocystis sp.* | Nitrate | 1.5 |
| F | 3N5L | Q9HYL8 | *Pseudomonas aeruginosa* | Phosphonate | 1.97 |
| F | 2O1M | O34852 | *Escherichia coli* | Arginine | 2.0 |
| F | 2I4B | Q55460 | *Synechocystis sp.* | Bicarbonate | 1.35 |
| F | 6EMN | O69061 | *Pseudomonas stutzeri* | Phosphite | 1.25 |
| F | 5O37 | O69052 | *Pseudomonas stutzeri* | Methylphosphonate | 1.37 |
| F | 5LQ1 | Q119I9 | *Trichodesmium erythraeum* | Methylphosphonate | 1.41 |
| F | 3P7I | Q1R3F7 | *Escherichia coli* | 2-aminoethyl phosphonate | 1.71 |
| F | 1D9Y | Q50964 | *Neisseria gonorrhoeae* | Iron(III) ion | 2.2 |
| F | 3I6V | Q5LTV6 | *Ruegeria pomeroyi* | Lysine | 2.0 |
| F | 3N26 | Q9Z869 | *Chlamydia pneumoniae* | Arginine | 2.1 |
| F | 5LQ5 | A3PC74 | *Prochlorococcus marinus* | Phosphite | 1.41 |
| F | 4PSH | Q9WZ62 | *Thermotoga maritima* | Arginine | 2.6 |
| F | 3KSX | Q8PHQ1 | *Xanthomonas citri* | Alkanesulfonate | 1.7 |
| F | 3UN6 | Q2G1I5 | *Staphylococcus aureus* | Zinc ion | 2.01 |
| F | 4YAH | P28635 | *Escherichia coli* | Methionine | 1.6 |
| F | 4LAT | Q97Q31 | *Streptococcus pneumoniae* | Phosphate | 1.88 |
| F | 3HV1 | Q5M4P7 | *Streptococcus thermophilus* | Polar amino acids | 1.9 |
| F | 2X26 | P75853 | *Escherichia coli* | Sulphonate | 1.75 |
| F | 2REG | Q92N37 | *Rhizobium meliloti* | Choline | 1.9 |
| F | 4GOT | O32167 | *Bacillus subtilis* | Methionine | 1.95 |
| F | 3TMG | O51169 | *Borrelia burgdorferi* | compatible solute Glycine<br>Betaine | 1.9 |
| F | 3IX1 | Q9K9G5 | *Bacillus halodurans* | N-formyl-4-amino-5-aminomethyl-2-methylpyrimidine | 2.4 |

## References

Ahmed AH, Wang S, Chuang HH, Oswald RE. 2011. Mechanism of AMPA receptor activation by partial agonists: disulfide trapping of closed lobe conformations. J Biol Chem 286:35257–35266. doi:10.1074/jbc.M111.269001

Armstrong Neali, Gouaux E. 2000. Mechanisms for Activation and Antagonism of an AMPA-Sensitive Glutamate Receptor. Neuron 28:165–181. doi:10.1016/S0896-6273(00)00094-5

Armstrong N, Gouaux E. 2000. Mechanisms for activation and antagonism of an AMPA-sensitive glutamate receptor: crystal structures of the GluR2 ligand binding core. Neuron 28:165–181. doi:10.1016/S0896-6273(00)00094-5

Armstrong N, Jasti J, Beich-Frandsen M, Gouaux E. 2006. Measurement of Conformational Changes accompanying Desensitization in an Ionotropic Glutamate Receptor. Cell 127:85–97. doi:10.1016/j.cell.2006.08.037

Berntsson RP a, Smits SHJ, Schmitt L, Slotboom DJ, Poolman B. 2010. A structural classification of substrate-binding proteins. FEBS Lett 584:2606–2617. doi:10.1016/j.febslet.2010.04.043

Boer M De, Gouridis G, Vietrov R, Begg SL, Schuurman-wolters GK, Husada F, Eleftheriadis N, Poolman B, Mcdevitt CA, Cordes T. 2019. Conformational and dynamic plasticity in substrate-binding proteins underlies selective transport in ABC importers. Elife 8:1–28. doi:10.7554/eLife.44652

Chen GQ, Cui C, Mayer ML, Gouaux E. 1999. Functional characterization of a potassium-selective prokaryotic glutamate receptor. Nature 402:817–821. doi:10.1038/45568

Doyle D a, Morais Cabral J, Pfuetzner R a, Kuo a, Gulbis JM, Cohen SL, Chait BT, MacKinnon R. 1998. The structure of the potassium channel: molecular basis of K+ conduction and selectivity. Science 280:69–77. doi:10.1126/science.280.5360.69

Dürr K, Chen L, Stein R, DeZorzi R, Folea IM, Walz T, Mchaourab H, Gouaux E. 2014. Structure and Dynamics of AMPA Receptor GluA2 in Resting, Pre-Open, and Desensitized States. Cell 158:778–792. doi:10.1016/j.cell.2014.07.023

Felder CB, Graul RC, Lee a Y, Merkle HP, Sadee W. 1999. The Venus flytrap of periplasmic binding proteins: an ancient protein module present in multiple drug receptors. AAPS PharmSci 1:7–26. doi:10.1208/ps010202

Felsenstein J. 2004. PHYLIP (Phylogeny Inference Package) version 3.6. Distributed by the author. http://www.Evolgswashingtonedu/phyliphtml.

Fleck MW, Cornell E, Mah SJ. 2003. Amino-acid residues involved in glutamate receptor 6 kainate receptor gating and desensitization. J Neurosci 23:1219–1227. doi:10.1523/jneurosci.23-04-01219.2003

Fukami-Kobayashi K, Tateno Y, Nishikawa K. 1999. Domain dislocation: a change of core structure in periplasmic binding proteins in their evolutionary history. J Mol Biol 286:279–290. doi:10.1006/jmbi.1998.2454

Gill A, Birdsey-benson A, Jones BL, Henderson LP, Madden DR. 2008. Correlating AMPA Receptor Activation and Cleft Closure across Subunits: Crystal Structures of the GluR4 Ligand-Binding Domain in Complex with Full and Partial 47:13831–13841. doi:10.1021/bi8013196

Gouridis G, Schuurman-wolters GK, Ploetz E, Husada F, Vietrov R, Boer M De, Cordes T, Poolman B. 2014. Conformational dynamics in substrate-binding domains influences transport in the ABC importer GlnPQ. Nat Publ Gr 22:57–64. doi:10.1038/nsmb.2929

Hanekop N, Höing M, Sohn-Bösser L, Jebbar M, Schmitt L, Bremer E. 2007. Crystal Structure of the Ligand-Binding Protein EhuB from Sinorhizobium meliloti Reveals Substrate Recognition of the Compatible Solutes Ectoine and Hydroxyectoine. J Mol Biol 374:1237–1250. doi:10.1016/j.jmb.2007.09.071

Horning MS, Mayer ML. 2004. Regulation of AMPA Receptor Gating by Ligand Binding Core Dimers. Neuron 41:379–388. doi:10.1016/S0896-6273(04)00018-2

Janovjak H, Sandoz G, Isacoff EY. 2011. A modern ionotropic glutamate receptor with a K(+) selectivity signature sequence. Nat Commun 2:232. doi:10.1038/ncomms1231

Krissinel E, Henrick K. 2004. Secondary-structure matching (SSM), a new tool for fast protein structure alignment in three dimensions. Acta Crystallogr Sect D Biol Crystallogr 60:2256–2268. doi:10.1107/S0907444904026460

Kuner T, Seeburg PH, Guy HR. 2003. A common architecture for K+ channels and ionotropic glutamate receptors? Trends Neurosci 26:27–32. doi:10.1016/S0166-2236(02)00010-3

L. Chen, K. Dürr EG. 2014. X-ray structures of AMPA receptor–cone snail toxin complexes illuminate activation mechanism. Science (80-) 345:1021–1026. doi:10.1126/science.1258409

Lam HM, Chiu J, Hsieh MH, Meisel L, Oliveira IC, Shin M, Coruzzi G. 1998. Glutamate-receptor genes in plants. Nature 396:125–126. doi:10.1038/24066

Lau AY, Roux B. 2011. The hidden energetics of ligand binding and activation in a glutamate receptor. Nat Struct Mol Biol 18:283–287. doi:10.1038/nsmb.2010

Lewis M, Chang G, Horton NC, Kercher M a, Pace HC, Schumacher M a, Brennan RG, Lu P. 1996. Crystal structure of the lactose operon repressor and its complexes with DNA and inducer. Science (80-) 271:1247–1254. doi:10.1126/science.271.5253.1247

Madej T, Lanczycki CJ, Zhang D, Thiessen P a., Geer RC, Marchler-Bauer A, Bryant SH. 2014. MMDB and VAST+: Tracking structural similarities between macromolecular complexes. Nucleic Acids Res 42:297–303. doi:10.1093/nar/gkt1208

Maltsev AS, Ahmed AH, Fenwick MK, Jane DE, Oswald RE. 2008. Mechanism of Partial Agonism at the GluR2 AMPA Receptor: Measurements of Lobe Orientation in Solution. Biochemistry 47:10600–10610. doi:10.1021/bi800843c

Mao B, Mccammon A. 1984. Structural Study of Hinge Bending in L-Arabinose-binding Protein. J Biol Chem 259:4964–4970.

Mayer ML, Olson R, Gouaux E. 2001. Mechanisms for ligand binding to GluR0 ion channels: crystal structures of the glutamate and serine complexes and a closed apo state. J Mol Biol 311:815–836. doi:10.1006/jmbi.2001.4884

Meyerson JR, Kumar J, Chittori S, Rao P, Pierson J, Bartesaghi A, Mayer ML, Subramaniam S. 2014. Structural mechanism of glutamate receptor activation and desensitization. Nature 57:328–334. doi:10.1038/nature13603

Nielsen H, Engelbrecht J, Brunak S, von Heijne G. 1997. Identification of prokaryotic and eukaryotic signal peptides and prediction of their cleavage sites. Protein Eng 10:1–6. doi:10.1142/S0129065797000537

Partin KM, Fleck MW, Mayer ML. 1996. AMPA receptor flip/flop mutants affecting deactivation, desensitization, and modulation by cyclothiazide, aniracetam, and thiocyanate. J Neurosci 16:6634–6647. doi:10.1523/jneurosci.16-21-06634.1996

Pettersen EF, Goddard TD, Huang CC, Couch GS, Greenblatt DM, Meng EC, Ferrin TE. 2004. UCSF Chimera - A visualization system for exploratory research and analysis. J Comput Chem 25:1605–1612. doi:10.1002/jcc.20084

Plested AJR. 2016. Structural mechanisms of activation and desensitization in neurotransmitter-gated ion channels. Nat Struct Mol Biol 23:494–502. doi:10.1038/nsmb.3214

Price MB, Jelesko J, Okumoto S. 2012. Glutamate Receptor Homologs in Plants: Functions and Evolutionary Origins. Front Plant Sci 3:1–10. doi:10.3389/fpls.2012.00235

Ramos-Vicente D, Ji J, Gratacòs-Batlle E, Gou G, Reig-Viader R, Luís J, Demian B, Navas-Perez E, García-Fernández J, Fuentes-Prior P, Escriva H, Roher N, Soto D, Bayés Á. 2018. Metazoan evolution of glutamate receptors reveals unreported phylogenetic groups and divergent lineage-specific events. Elife 7:1–36. doi:10.7554/eLife.35774

Salazar H, Eibl C, Chebli M, Plested A. 2017. Mechanism of partial agonism in AMPA-type glutamate receptors. Nat Commun 8:1–11. doi:10.1038/ncomms14327

Schauder DM, Kuybeda O, Zhang J, Klymko K, Bartesaghi A, Borgnia MJ, Mayer ML, Subramaniam S. 2013. Glutamate receptor desensitization is mediated by changes in quaternary structure of the ligand binding domain. Proc Natl Acad Sci U S A 110:5921–6. doi:10.1073/pnas.1217549110

Scheepers GH, Lycklama a Nijeholt JA, Poolman B. 2016. An updated structural classification of substrate-binding proteins. FEBS Lett 590:4393–4401. doi:10.1002/1873-3468.12445

Schönrock M, Thiel G, Laube B. 2019. Coupling of a viral K+-channel with a glutamate-binding-domain highlights the modular design of ionotropic glutamate-receptors. Commun Biol 2. doi:10.1038/s42003-019-0320-y

Sobolevsky AI, Rosconi MP, Gouaux E. 2009a. X-ray structure, symmetry and mechanism of an AMPA-subtype glutamate receptor. Nature 462:745–756. doi:10.1038/nature08624

Sobolevsky AI, Rosconi MP, Gouaux E. 2009b. X-ray structure, symmetry and mechanism of an AMPA-subtype glutamate receptor. Nature 462:745–756. doi:10.1038/nature08624

Sobolevsky AI, Rosconi MP, Gouaux E. 2009c. X-ray structure, symmetry and mechanism of an AMPA-subtype glutamate receptor. Nature 462. doi:10.1038/nature08624

Sun Y, Olson R, Horning M, Armstrong N, Mayer M, Gouaux E. 2002. Mechanism of glutamate receptor desensitization. Nature 417:245–253. doi:10.1038/417245a

Tam R, Saier MH. 1993. Structural, Functional, and Evolutionary Relationships among Extracellular Solute-Binding Receptors of Bacteria. Microbiol Rev 57:320–346.

Tang C, Schwieters CD, Clore GM. 2007. Open-to-closed transition in apo maltose-binding protein observed by paramagnetic NMR 449:1078–1082. doi:10.1038/nature06232

Tikhonov DB, Magazanik LG. 2009. Origin and molecular evolution of ionotropic glutamate receptors. Neurosci Behav Physiol 39:763–773. doi:10.1007/s11055-009-9195-6

Traynelis SF, Wollmuth LP, McBain CJ, Menniti FS, Vance KM, Ogden KK, Hansen KB, Yuan H, Myers SJ, Dingledine R. 2010. Glutamate Receptor Ion Channels: Structure, Regulation, and Function. Pharmacol Rev 62:405–496. doi:10.1124/pr.109.002451

Twomey EC, Sobolevsky AI. 2018. Structural Mechanisms of Gating in Ionotropic Glutamate Receptors. Biochemistry 57:267–276. doi:10.1021/acs.biochem.7b00891

Weston MC, Schuck P, Ghosal A, Rosenmund C, Mayer ML. 2006. Conformational restriction blocks glutamate receptor desensitization. Nat Struct Mol Biol 13:1120–1127. doi:10.1038/nsmb1178

Wood MW, Vandongen HMA, Vandongen AMJ. 1995. Structural conservation of ion conduction pathways in K channels and glutamate receptors. Proc Natl Acad Sci USA 92:4882–4886.

Yelshanskaya M V., Li M, Sobolevsky a. I. 2014. Structure of an agonist-bound ionotropic glutamate receptor. Science (80-) 345:1070–1074. doi:10.1126/science.1256508

Zhang W, Cho Y, Lolis E, Howe JR. 2008. Structural and Single-Channel Results Indicate That the Rates of Ligand Binding Domain Closing and Opening Directly Impact AMPA Receptor Gating 28:932–943. doi:10.1523/JNEUROSCI.3309-07.2008

